# Neural auditory contrast enhancement in humans

**DOI:** 10.1101/458521

**Authors:** Anahita H. Mehta, Lei Feng, Andrew J. Oxenham

## Abstract

The perception of sensory events can be suppressed or enhanced by the surrounding spatial and temporal context in ways that help in detecting novel objects and establishing perceptual constancy. In the auditory system, the phenomenon known as auditory enhancement reflects a general principle of contrast enhancement, where a target sound embedded within a background sound becomes perceptually more salient if the background is presented first by itself. This effect is highly robust, producing an effective enhancement of the target of between 5 and 25 dB (more than two orders of magnitude in intensity), depending on the task. Despite their ubiquity in vision, neural correlates of auditory contrast enhancement have yet to be identified in humans. Here we used the auditory steady-state response to probe the neural response to a target sound under conditions of enhancement. The probe was simultaneously modulated in amplitude with two modulation frequencies, to distinguish cortical from subcortical contributions to this phenomenon. We found robust auditory cortical, but not subcortical, enhancement that correlates with behavior and is consistent with an early theoretical model that postulates neural adaptation of inhibition. Our findings provide empirical support for a previously unverified theory of auditory enhancement and point to new approaches for improving sensory prostheses for hearing loss, such as hearing aids and cochlear implants.

**Significance Statement:** A target sound embedded within a background sound becomes perceptually more salient if the background is presented first by itself. This phenomenon, where the target “pops out”, is known as auditory enhancement. It reflects a general principle of contrast enhancement, and helps in the detection of new acoustic events in the environment and in establishing the perceptual constancy of speech and other biologically relevant sounds under varying acoustic conditions. We use EEG in humans to reveal a cortical correlate of this perceptual phenomenon that provides empirical support for a longstanding but previously unverified theoretical account.

## Introduction

Across sensory domains, perception and its underlying neural substrates are highly dependent on context (1). Such context dependence improves neural coding efficiency (2, 3) and plays a key role in adapting sensory systems to the long-term properties of the environment, thereby enhancing sensitivity to novel events (4) and helping to establish perceptual constancy under a wide range of backgrounds and environments (5). Context effects in hearing can be illustrated with an effect known as auditory enhancement (6, 7). In this paradigm, a target embedded within a background sound can “pop out” perceptually if the background sound is presented by itself first (Figure 1A). This effect is highly robust, with the enhancement of the target corresponding to an intensity increase of between 5 and 25 dB (i.e., more than two orders of magnitude), depending on the task (8, 9), and is also ecologically relevant, as it reflects properties that strongly affect speech perception (10–12).

**Figure 1:**
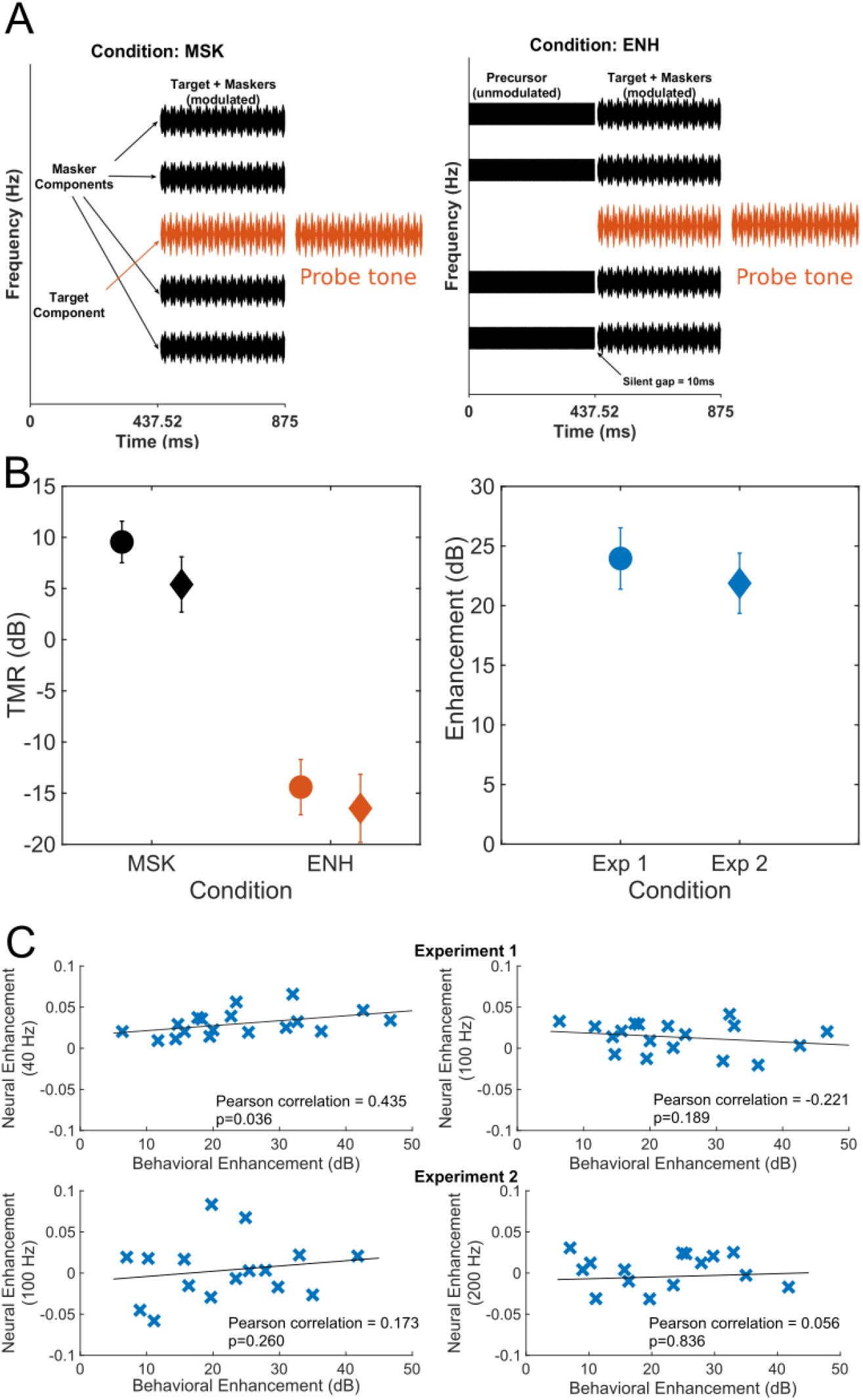
**A.** Schematic representation of the stimuli used for measuring behavioral thresholds. **B.** Left panel shows average thresholds with and without precursors (ENH and MSK respectively) for experiments 1 (N=18) and 2 (N=16). Right panel shows the amount of enhancement, defined as the difference in the MSK and ENH thresholds, for each experiment. The circles indicate mean thresholds for experiment 1. The diamonds indicate mean thresholds for experiment 2. Error bars represent ±1 standard error of the mean. **C.** Correlations between neural and behavioral enhancement at low and high frequencies for experiments 1 and 2. The neural enhancement for experiment 1 was calculated as the average PLV enhancement across the three TMRs.

Despite its central role in auditory perception, little is known about the neural mechanisms underlying auditory enhancement. One proposal, known as adaptation of inhibition (6), invokes lateral inhibition between adjacent spectral components (13), comparable to the well-known center-surround receptive fields in the visual system (14), but with the inhibition adapting over time. Thus, when a new spectral component is introduced to an ongoing stimulus, the inhibition from the surrounding components has adapted, leading to an enhanced representation of the new component. Although this theoretical framework can account for a variety of behavioral results (6–8, 15–19), little evidence exists for its neural implementation. Physiological studies in non-human species have found no evidence of enhancement at the level of the auditory nerve (20) but have reported some neural responses consistent with adaptation of inhibition at the level of the inferior colliculus (IC) (21). Taken together, the results suggest hierarchical processing, with enhancement only emerging at the level of the IC. However, such an interpretation must be tempered by the fact that the studies were carried out in different species, only one study was attempted in an awake preparation (21) and none of the studies included behavior. Evidence from human studies remains limited. No evidence for enhancement has been found in the cochlea, using otoacoustic emissions (OAEs) (22) or in auditory neural pathways using the 80-Hz auditory steady-state responses (ASSR) (23). Thus, no neural correlates of enhancement have yet been reported in humans.

In the current study, we introduce a method of targeting both subcortical and cortical auditory structures in humans using ASSR measured with electroencephalography (EEG). We combine a stimulus configuration that produces up to 24 dB of behavioral enhancement (8) with a frequency-tagging paradigm, where the neural responses to both the masker and target components can be teased apart based on differences in the modulation rate of the envelopes applied to the target and masker. In experiment 1, we applied amplitude modulation (AM) to each component with two rates to simultaneously record responses, quantified as phase locking values (PLVs) (24), from slower (~40 Hz) and faster (~100 Hz) rates that are thought to target primarily cortical and subcortical levels of the auditory pathways, respectively. In experiment 2, we focused on subcortical responses by using only a single high modulation rate, around either 100 or 200 Hz. We also measured the behavioral thresholds for the stimuli used in the EEG experiments in all participants to ensure that the stimuli evoked perceptual enhancement (Figure 1B). Overall, we observed robust correlates of perceptual enhancement for the low frequencies (~40 Hz), thought to reflect cortical responses (25–28), but found little or no evidence for enhancement at the higher frequencies (~100 and 200 Hz), emanating from primarily subcortical sources (29–31). The pattern of cortical responses were consistent with predictions based on the longstanding, but previously unverified, theory of enhancement based on adaptation of neural inhibition.

## RESULTS

### Behavioral thresholds

We measured the behavioral detection thresholds for the target, using the same stimuli that were used in both the EEG experiments in all participants to ensure that the stimuli evoked perceptual enhancement (Figure 1A). Threshold was defined as the minimum target level required to correctly judge whether the subsequent probe was presented at the same frequency as the target. The mean behavioral thresholds for Experiments 1 and 2 are shown in Fig. 1B (see Supplementary Figure S1 for individual thresholds). The mean target-to masker ratio (TMR) in Experiment 1 was 9.5 dB in the MSK condition and −14.4 dB in the ENH condition. The mean TMR in Experiment 2 was 5.4 dB TMR in the MSK condition and −16.5 dB in the ENH condition. Thus, the average amount of enhancement calculated as the difference in thresholds with and without the precursors (MSK - ENH) was 23.9 dB for Experiment 1 and 21.9 for Experiment 2. This value is comparable to the average enhancement reported in an earlier study with similar stimuli (8). A repeated-measures analysis of variance (ANOVA) carried out on the thresholds indicated a significant effect of masking condition, with and without the precursor (MSK vs ENH) (*F*_1,15_ = 182.6; *p* <0.0005, η_p_^2^ =0.924), but no significant effect of experiment (Experiment 1 vs. 2) (*F*_1,15_ = 2.82; *p* = 0.114, η_p_^2^ =0.158), and no significant interaction between the two factors (*F*_1,15_ = 0.08; *p* = 0.782, η_p_^2^ =0.005), confirming the presence of enhancement, and suggesting that the different amplitude modulations used for the two EEG experiments did not affect the amount of enhancement measured behaviorally.

### EEG data

Data from a representative participant in Experiment 1 are shown in Fig. 2B. There are four distinct peaks in PLV at the four tagging frequencies (two for the maskers and two for the target). Figure 2C shows the average PLVs at the target and masker modulation frequencies from the 18 participants in Experiment 1 as a function of the TMR for the lower and higher modulation frequencies (left and right panels, respectively). We observed robustly higher PLVs at the target frequency of 43 Hz in the ENH condition than in the MSK condition, indicating enhancement, but no change in the PLVs at the masker frequency of 34 Hz (Figure 2C, left panels). At the higher frequencies (around 100 Hz), the amount of target enhancement was less (Figure 2C, right panels). A repeated-measures ANOVA with the PLV as the dependent variable and within-subjects factors of modulation rate (~40 Hz or ~100 Hz), stimulus component (target or masker), presence of precursor (MSK or ENH), and TMR (0, −5, or −10 dB), showed that all main effects were significant (precursor: F_1,17_=12.12, p=0.003, η_p_^2^ =0.416; stimulus component: F_1,17_=5.02, p=0.039, η_p_^2^ =0.228; modulation rate: F_1,17_=5.33, p=0.034, η_p_^2^ =0.416; TMR: F_2,34_=3.31, p=0.049, η_p_^2^ =0.163). Significant interactions were observed between presence of precursor and stimulus component (F_1,17_=47.55, p<0.001, η_p_^2^ =0.737), presence of precursor and modulation rate (F_1,17_=12.15; p=0.003, η_p_^2^ =0.417), and stimulus component and TMR (F_2,34_ = 38.28, p<0.001, η_p_^2^ =0.692). The three-way interaction was also significant (F_1,17_=11.52; p=0.003, η_p_^2^ =0.404). No other interactions reached significance (p > 0.05 in all cases). Post hoc comparisons (with Bonferroni correction for multiple comparisons) showed that PLVs were significantly different between the MSK and ENH conditions only for the target components (for the low rate, p<0.001; for the high rate, p=0.034). Overall enhancement, defined as the difference between the target PLV in the ENH and MSK conditions, subtracted by the difference in the masker PLV in the ENH and MSK conditions (Methods, Eq 3), was significantly greater than zero (Figure 2C bottom panels) at all three TMRs for the low (~40 Hz) rate (TMR0: t17=3.3,p=0.004, TMR-5: t17=6.3, p<0.001, TMR-10: t17=4.33, p<0.001) but was only significantly greater than zero at −5 dB TMR for the high (~100 Hz) rate (TMR0: t17=1.12,p=0.263, TMR-5: t17=3.8, p=0.001, TMR-10: t17=1.46, p=0.161). The pattern of results, especially at the lower modulation frequency (~40 Hz), is consistent with predictions based on the theory of adaptation of inhibition, as outlined in the Introduction: the precursor should result in an increase in the response to the target, but no decrease in the response to the masker.

**Figure 2:**
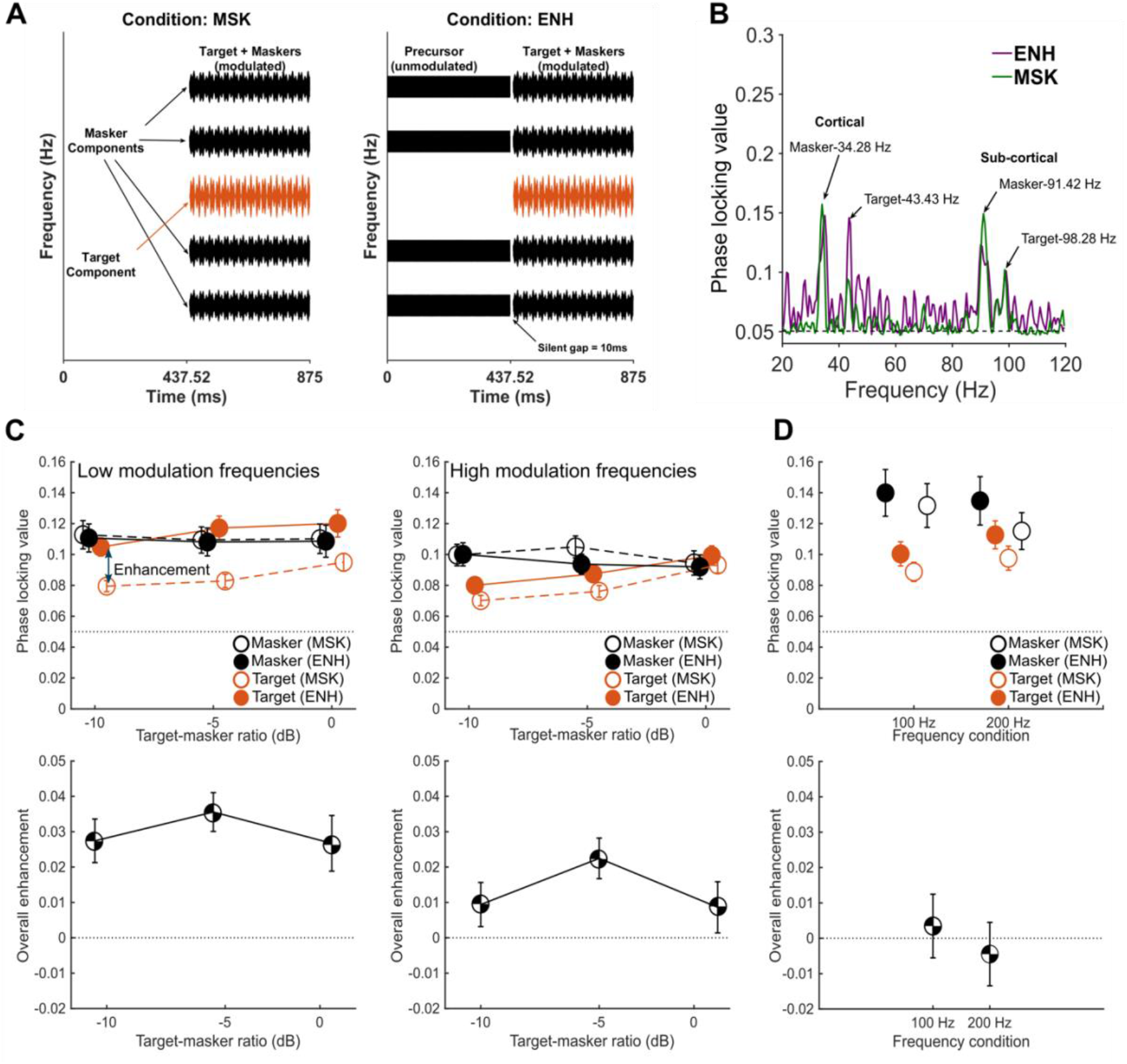
**A.** Schematic diagram of the stimuli with amplitude-modulated tones used in the EEG measurements for Experiments 1 and 2. **B**. Examples of phase locking values (PLVs) from one participant in Experiment 1. PLVs are plotted as a function of frequency measured at TMR −5 in ENH (purple) and MSK (green) conditions. Each curve represents the averaged PLVs across 30 electrodes. The arrows point to the four distinct peaks at all four tagging frequencies used for the maskers and target. Note that the amplitude of the PLV for the masker components remain similar in both conditions (MSK and ENH), whereas the target component amplitudes are enhanced in the ENH conditions. The dashed line indicates the noise floor. **C.** (Top row) PLVs across all 18 participants for both low (left) and high (right) tagging frequencies for all components across TMRs in Experiment 1. (Bottom row) Average overall enhancement, defined as the difference in the enhancement of the PLV for the targets and the maskers (see Eq. 3 in Methods), for both low (left) and high (right) tagging frequencies. **D.** (Top row) PLVs across all 16 participants for both sets of tagging frequencies for all components in Experiment 2. (Bottom row) Average overall enhancement, defined as the difference in the enhancement of the PLV for the targets and the maskers (see Eq. 3 in Methods), for both frequency conditions. All error bars in Figure 2 represent ±1 standard error of the mean. The dashed lines in the top rows of panels C and D indicate the noise floor. The dashed lines in the bottom rows of panels C and D indicate no enhancement.

The results from the higher modulation frequency (~100 Hz) are more difficult to interpret for two reasons. First, the observed amount of enhancement is less robust and was only statistically significant at one of the three TMRs; second, the source of the ASSRs around 100 Hz remains somewhat in dispute, with some claiming a significant cortical contribution (29, 30) and others finding the response dominated by subcortical structures (29, 30).

There is more general agreement that the sources of responses at higher modulation frequencies (~200 Hz) are subcortical, because of the limited ability of the auditory cortex to synchronize to high-frequency stimuli (30, 31). For this reason, we ran a second experiment to test both 100- and 200-Hz modulation rates, using only single modulation rate on each carrier, so that the modulation depth could be increased to 100% for the target, thereby increasing the overall neural response to the modulation (34). Only the TMR that had initially indicated possible enhancement at 100 Hz was included (−5 dB). The resulting PLVs were well above the noise floor (Figure 2D upper panel), even exceeding those measured for the lower-frequency (cortical) responses in Experiment 1. However, no clear enhancement effects were observed at either the 100- or 200-Hz rate (Figure 2D lower panel). A repeated-measures ANOVA with PLV as the dependent variable and within-subjects factors of modulation rate (100 or 200 Hz), stimulus component (target or masker), and presence of precursor (MSK or ENH) revealed main effects of precursor (F_1,15_=10.43, p=0.006, η_p_^2^ =0.410) and stimulus component (F_1,15_=17.05, p=0.001, η_p_^2^ =0.532), but no main effect modulation rate (F_1,15_=0.002, p=0.987, η_p_^2^ =0.000). No interactions were significant (p > 0.05 in all cases), suggesting that effects of the precursor did not differ between the target and masker. In line with this outcome, overall enhancement (effect of precursor on the target PLV subtracted by its effect on the masker PLV) was not significantly different from zero (100 Hz: t15=0.37, p=0.72; 200 Hz: t15=-0.48, p=0.64). Overall, our results provide no evidence for a subcortical contribution to auditory enhancement. A complete table of statistical values for all comparisons in both experiments can be found in Supplementary Tables 1 and 2.

### Correlation between EEG and behavior

We explored a possible relationship between neural responses and behavior at the level of individual participants by correlating the overall PLV enhancement measure from each participant, averaged across TMRs, with their behavioral detection thresholds using the same stimulus configurations. We found a small but significant positive behavioral correlation between the neural and behavioral enhancement at the low modulation rates (Figure 1C, top left panel; Pearson correlation, r = 0.435, p=0.036, one tailed). We found no significant correlations at the higher modulation rates (around 100 or 200 Hz) in either the first or second experiment (Figure 1C, top right panel and bottom row).

## DISCUSSION

The current study reveals a neural correlate of auditory enhancement in humans. This correlate was observed at cortical but not sub-cortical levels. Importantly, the enhancement is produced by an increase in the response to the target, rather than an adaptation or decrease in the response to the maskers, in line with an early theory of auditory enhancement, known as adaptation of inhibition (6).

### Neural gain and behavioral thresholds

A previous study found no evidence of neural enhancement in the 80-Hz ASSRs (23). The lack of neural enhancement observed at our higher modulation frequencies is in line with those earlier findings. However, our paradigm also yields perceptual enhancement of 20-25 dB, compared to about 5 dB enhancement in the earlier study, making our findings less likely to be affected by factors related to a smaller behavioral effect size. Part of the difference in behavioral outcomes between the two paradigms may be due to our use of frequency roving from trial to trial, which reduces the possibility of contamination via longer-term adaptation effects between trials (8).

The enhanced neural responses to the target in the presence of the precursor seen at ~40 Hz in our current study correlate with the degree of perceptual enhancement (Figure 1C). However, the precise relationship between the neural representations and perception of the target remains unclear. For instance, the average behavioral threshold in the MSK condition is around 10 dB TMR (Fig. 1B), whereas a robust neural representation of the target can be observed at TMR values well below that (e.g., at −10 dB TMR), suggesting that a cortical representation of the target that persists well below its perceptual threshold. Nevertheless, assuming a constant relationship between the behavioral audibility of the target and its PLV, the amount of perceptual auditory enhancement (in dB) should be reflected in the difference of TMRs that yield the same target PLV in the MSK and ENH conditions (orange lines in Fig. 2C). For instance, the average PLV of the cortical response (left panel in Fig. 2C) at −10 dB TMR in the ENH condition is greater than the PLV in the MSK condition at 0 dB TMR. In this case, the cortical responses would predict perceptual enhancement greater than 10 dB, in line with the behavioral results indicating enhancement of 20 dB or more.

### Possible neural mechanisms of auditory enhancement

A possible neural mechanism underlying auditory enhancement is the adaptation of inhibition (6). In the central auditory system, starting from the cochlear nucleus, across-frequency processing starts to emerge, where neurons selective to one center frequency can be laterally suppressed or inhibited by neighboring frequencies (35, 36). When a complex tone is presented, the neurons that respond to each component also mutually inhibit each other. Since the frequency-specific responses are known to adapt over time (1, 37), it is possible that this form of inhibition adapts in a similar way. In the ENH condition, the inhibition to the target response from the masker may be adapted by the precursor, resulting in an absolute increase in the target response. Previous single-unit studies have found evidence for this mechanism by showing increased neural firings in the inferior colliculus (21). Our results with the 40-Hz ASSR are consistent with the predictions of this mechanism: the precursor produces no decrease in the response to the maskers (black lines in Figure 2C), suggesting no net adaptation of the masker components, and instead produces an increase in the response to the target, as would be expected if the inhibition of the target had been adapted by the precursor.

This lack of adaptation in the masker responses could also be explained by the adaptation of inhibition mechanism: since neurons responding to the four maskers are laterally inhibited by each other, the responses to the maskers should decrease due to adaptation from the precursor, but this adaptation may be counteracted by the adaptation of the lateral inhibition, leading to no net change in responses.

### Differential contributions to the frequency following response

The double-modulation paradigm used in this study provides us with the ability to parse out the contributions from various stages of the auditory pathway simultaneously. Although there is broad consensus that the ASSR around 40 Hz arises primarily from auditory cortex, with only a weak contribution from subcortical structures (25–28), there remains debate surrounding the primary source of the ASSR around 100 Hz (26–31, 38, 39).

Cortical contributions have been shown to dominated the ASSR or frequency following response (FFR) to the voice pitch (F0 = 100 Hz) of speech, as measured with MEG (32), and a correlation has been reported between the strength of the FFR to F0 in EEG and BOLD signal in the right posterior auditory cortex using fMRI (33). However, there is a natural bias of MEG toward superficial brain tissue (40), and the relation between the BOLD signal in fMRI and underlying neural activity still remains an open question (41). Indeed, another study (29) showed that subcortical structures (auditory nerves and brainstem) make the largest contribution to FFRs recorded using EEG and found that cortical sources contributed little to the FFR above 100 Hz.

Given the lack of consensus on the source of responses to 100 Hz, we extended our modulation frequencies to 200 Hz, where there is general agreement that neural measures reflect responses from primarily subcortical structures (30, 31). Based on the fact that no significant enhancement was observed at either frequency in Experiment 2, we can conclude that our method has revealed cortical, but not subcortical, correlates of auditory enhancement.

### Inherited or emergent in auditory cortex?

In the current study, the enhanced neural responses to the target were reflected in the 40-Hz ASSRs but not in the 100- or 200-Hz ASSRs. Although we found no evidence for enhancement at the higher modulation frequencies, we cannot rule out its existence based on our surface EEG measurements. For instance, it is possible that the enhanced target responses emerge in the inferior colliculus, as suggested by the enhanced firing rate of single neurons (21), and the increased amplitude of 100-Hz and 200-Hz ASSRs in some of the participants (see Supplementary Figures S3 and S4). However, evidence for this activity is not reliably detected by surface EEG electrodes. This could be because neurons in subcortical structures that reflect enhancement may not effectively entrain to the stimulus envelope, as required by our method. Alternatively, enhancement could have a cortical origin and the neural enhancement observed in the brainstem could come from descending (efferent) corticofugal projections (42), which would be less likely to be entrained. Future studies using animal models using the behavioral paradigm employed here may shed more light on the locus of this phenomenon and answer the question of whether it is achieved via feedforward or feedback mechanisms originating in the cortex, by examining the neural responses in the brainstem when corticofugal neurons are deactivated by cooling, optogenetic silencing, or other pharmaceutical manipulations.

## METHODS

### Participants

Eighteen participants (11 female and 7 male) took part in Experiment 1 and 16 (11 females and 5 males) in Experiment 2. The participants were between 18 and 34 years old, had normal hearing, as defined by audiometric pure-tone thresholds better than 20 dB hearing level (HL) in both ears at octave frequencies from 250 to 8000 Hz, and had no reported history of hearing or neurological disorders. All participants provided written informed consent and were compensated for their time. All protocols were approved by the University of Minnesota Institutional Review Board.

### Behavioral measures of enhancement

Perceptual thresholds for a target tone were measured in conditions both with and without a precursor stimulus, in order to confirm and quantify the amount of behavioral enhancement. The same stimuli were then employed in the EEG recordings, similar to the task used in a previous study (8).

#### Stimuli

A schematic representation of the behavioral stimuli is shown in Figure 1A. In the simultaneous masker condition with no precursor (MSK), each trial contained an inharmonic complex tone with five equal-amplitude components spaced apart from each other by 5/11 octaves, followed by a pure-tone probe. The base frequency for the conditions with modulation frequencies around 100 Hz or lower was 602 Hz (in both experiment 1 and 2) and was 1204 Hz for conditions with modulation frequencies around 200 Hz. The target tone was the 3^rd^ component within the complex. The frequency of the probe tone was either the same as the frequency of the target tone, or was geometrically centered between the frequencies of the target tone and of one of its adjacent neighbors (either the lower or the upper neighbor). From trial to trial, all frequencies in the inharmonic complex were randomly roved together, with uniform distribution, on a logarithmic scale. The amount of roving was either within a one-octave frequency range (experiment 1) or a half-octave frequency range (experiment 2) centered on the base frequencies. The inharmonic complex and probe tone were each 437.52 ms long, including 10-ms raised-cosine onset and offset ramps, separated by a 100-ms silent gap. The level of each masker component was 45 dB sound pressure level (SPL). The level of the target tone was adaptively varied, as described in the Procedure section below. In the enhanced condition (ENH), a precursor was presented before the inharmonic complex. The four precursor frequencies matched those of the masker in each trial (i.e., no component at the target frequency). The precursor components were presented at the same level as the masker components. The duration of the precursor was also 437.52 ms, including 10-ms raised-cosine onset and offset ramps. The delay between the precursor offset and inharmonic complex onset was 10 ms. In Experiment 1, the four masker components in the masker-plus-target complex were amplitude modulated with the sum of two sinusoidal waveforms at 34.28 and 91.42 Hz, each presented at a modulation depth of 25%. The amplitude modulation for the target component was the sum of two other sinusoidal waveforms at 43.43 and 98.28 Hz, each modulated at a depth of 50% (Figure 2A). The probe tone was modulated in the same way as the target. The 437.52-ms duration of the inharmonic complex and probe tone ensured an integer number of cycles of all the modulation frequencies, such that the starting and ending phases were zero for all modulators. The precursor components were not modulated in the ENH condition. In Experiment 2, the four masker components in the masker-plus-target complex were amplitude modulated with sinusoidal waveforms at 91.42 Hz for the 100-Hz conditions and 217.13 Hz for the 200-Hz condition, each presented at a modulation depth of 50%. The target component was amplitude modulated with sinusoidal waveforms at 98.28 Hz for the 100-Hz conditions and 233.13 Hz for the 200-Hz condition at a modulation depth of 100%.

#### Procedure

Participants were individually seated in a double-walled sound-attenuating booth. The stimuli were generated digitally using the AFC software package (43) under Matlab (Mathworks, Natick, MA) at a 48-kHz sampling rate, delivered through an E22 soundcard (LynxStudio, Costa Mesa, CA) with a 24-bit resolution, and presented monaurally to the right ear via HD650 headphones (Sennheiser, Old Lyme, CT). The task was a present/absent task where the listeners were asked to report whether or not the probe tone had been present in the target-plus-masker complex. The basic premise behind this task is that when the target tone is perceived in the presence of the maskers, it is possible to compare the pitch of the target tone and the probe tone. Hence, if the target tone is perceived, the listeners can then make a decision of whether the probe tone is the same frequency as the target tone. The two alternatives (probe tone frequency present or absent) were presented with equal *a priori* probability. For example, when the probe tone frequency matched the target tone frequency, a correct response would be ‘present’ whereas when the probe tone frequency was different from the target tone, the correct answer would be ‘absent’. When the probe-tone frequency was different, it was above or below the frequency of the target with equal a priori probability. The level of the target tone was initially set to 65 dB SPL (i.e., 20 dB higher than the individual masker components) and was varied adaptively following a two-down one-up rule that tracks the 70.7% correct point on the psychometric function (44). Feedback was provided after each trial. The level of the probe tone was always the same as that of the target. Initially the level of the target tone was varied in steps of 5 dB. After two reversals in the direction of the adaptive tracking procedure, the step size was reduced to 2 dB. The run was terminated after eight reversals and the threshold was computed as the average target level at the last six reversal points of the tracking procedure. There were four conditions in total, including a simultaneous masker with no precursor condition (MSK) and an enhanced condition (ENH), either with pure tones or AM tones. Each condition was tested twice for each participant in experiment 1 and four times in experiment 2. Each participant either started with the pure tones or the AM tones, with the order counterbalanced between participants. The presentation order of the MSK and ENH conditions were randomized separately for each participant and tone type (pure or AM). Threshold was defined in terms of the target-to-masker ratio (TMR), or the level of the target relative to the level per masker component.

#### Screening

To be eligible to take part in the EEG experiments, participants were required to pass the behavioral screening tests. As the behavioral task is based on pitch discrimination, the first screening test was comprised of two simple pitch discrimination tasks to screen out people with extremely poor pitch discrimination. In the first task, participants were presented with two consecutive pure tones, each 437.52 ms in duration, separated from each other by a silent gap of 100 ms. The two tones were either the same or differed in frequency by the same amount as the target and probe tones in the main experiment. Participants were asked whether the two tones had the same or different pitches. The task had 20 trials, 10 trials of which were the same pitch and 10 trials of which were different (higher or lower with equal a priori probability). In the second task, both tones were amplitude modulated with a sum of two sinusoids at 43.43 and 98.28 Hz with a 50% modulation depth for either frequency. The tones in the first task and the carriers of the second task were roved in frequency from trial to trial in the same way as in the main experiment. All participants had to obtain at least 80% correct in both sessions to pass. All participants for both experiments passed both the screening tasks. Once participants passed the pitch discrimination screening, the behavioral blocks to measure enhancement thresholds were administered (as described in the Stimuli and Procedure sections above). The screening criterion for recruitment for the EEG tasks was to obtain a threshold for the enhanced conditions below 0 dB TMR (see Supplementary Figure S1 for individual thresholds). For Experiment 1, 18 out of 20 recruited participants passed the criterion of having enhanced thresholds below 0 dB TMR whereas for Experiment 2, 16 out of 22 recruited participants passed this criterion.

### EEG measures of enhancement

#### Procedure

Each participant took part in one experimental session of 2.5 hours, including setup and EEG data collection. Participants were seated in a double-walled, electrically shielded, sound-attenuating booth. Before EEG data acquisition, participants were fitted with a cap containing 64 pin type active electrodes compatible with the BioSemi headcaps. Two additional reference electrodes, one placed on each mastoid, and two ocular electrodes, above and below the right eye, were used. It was ensured that the magnitude of the offset voltages for all electrodes was always < ±10 mV. The EEG data were recorded at a sampling rate of 4096 Hz using a 64-channel BioSemi Active Two EEG system (Amsterdam, Netherlands). The sounds were presented via ER-1 insert phones (Etymotic Research, Elk Grove, IL), and participants watched a silent movie with subtitles during data acquisition. The presentation order of the conditions was pseudorandomized across participants.

#### Experiment 1

In this experiment, we recorded the ASSR to estimate the population neural responses to the masker tones and target tone separately by tagging them with different signature AM frequencies. Each tone was modulated by one low rate (around 40 Hz) and one high rate (around 100 Hz). The stimuli used for the EEG experiment were the same AM tones used in the behavioral experiments, but without the probe tone (Figure 2A), which was not required as no behavioral thresholds were measured during the EEG sessions. Both conditions, with a precursor (ENH) and without a precursor (MSK), were tested at three target-to-masker ratios (TMRs) of 0, −5 and −10 dB, resulting in a total of six conditions. A total of 1000 trials were run in each condition for each participant, and the frequencies of the entire inharmonic complex were randomly roved within an octave on each presentation in the same way as for the behavioral measures in Experiment 1. Half the trials were presented in the inverted starting polarity. The durations of the precursor and masker were both 437.52 ms, including 10-ms raised-cosine onset and offset ramps in the same way as in the behavioral experiments.

#### Experiment 2

In this experiment we recorded the ASSR to estimate the population neural responses to the masker tones and target tone separately by tagging them with only one frequency per tone and testing frequencies either around 100 Hz or around 200 Hz. This experiment was also designed to elicit the maximum possible ASSR to the higher frequencies (around 100 and 200 Hz) to avoid the problems of the signal being too small to detect reliably above the noise floor. The stimuli used for this experiment were similar to those of Experiment 1 with the following changes: Both conditions, with precursor (ENH) and without precursor (MSK), were tested at both AM rates, but at only one target-to-masker ratio (TMRs) of −5 dB, resulting in a total of four conditions. A total of 1500 trials were run in each condition for each participant, and the frequencies of the entire inharmonic complex were randomly roved within half an octave on each presentation. The carrier frequency of the target ranged between approximately 1785 Hz and 2525 Hz for the 100-Hz AM conditions and between approximately 3570 Hz and 5050 Hz for the 200-Hz AM conditions. For the 100-Hz conditions, all four masker components in the masker-plus-target complex were amplitude modulated at 91.42 Hz with 50% modulation depth and the target component was modulated at 98.28 Hz with 100% modulation depth. For the 200-Hz conditions, all four masker components in the masker-plus-target complex were amplitude modulated at 217.13 Hz with 50% modulation depth and the target component was modulated at 233.13 Hz with 100% modulation depth. The precursor components in the ENH conditions were not modulated.

### EEG data analysis

The EEG pre-processing and averaging was done using the EEGLAB toolbox (45) for MATLAB (v14.1.1). The raw waveforms were down-sampled to 1024 Hz and re-referenced to the average of the two mastoids. The signals were bandpass filtered using a zero phase-shift FIR filter (from 1 to 100 Hz for Experiment 1 and from 1 to 250 Hz for experiment 2). For each condition, the continuous EEG time series was divided into epochs. For the MSK conditions, the epoch extended from 100 ms before stimulus onset to 700 ms post stimulus onset. For the ENH condition, the epoch extended from 100 ms pre-stimulus onset to 1100 ms post stimulus onset, since the stimulus was longer with the presence of the precursor. The EEG epoched signal was then baseline corrected relative to the 100-ms pre-stimulus baseline. Independent Component Analysis (ICA) was used to remove artefacts related to eye movements and blinks.

Further analysis was done in MATLAB. The discrete Fourier transforms (DFTs) of the processed EEG signals were applied to the time-domain waveforms from individual trials and the phases at each frequency were extracted. For each electrode in each condition for each participant, the PLV to the envelope was computed by averaging the phases of the individual trials’ responses at each frequency from 400 random samples (drawn with replacement) (24). The average phases were calculated for the 200 positive polarity trials (POSi) and 200 negative polarity trials (NEGi) separately beforehand (Eq. 1).

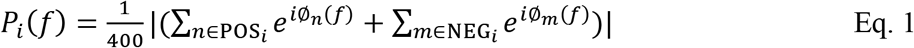

The same procedure was repeated 100 times independently to estimate the distribution of the PLVs and the mean was calculated as the observed PLVs for one electrode in one condition (Eq. 2).

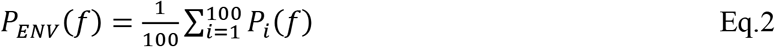

To evaluate the statistical significance of PLV values, bootstrapping was used to estimate the noise floor. A null model was tested by generating one random distribution of PLVs by repeating the procedure for PLV calculation described above 1000 times, except that the phase in Eq. 1 was set to be random (uniformly distributed from 0 to 2π, independently selected for each trial and repetition, *i).* The calculated mean PLV distributions from the experimental data can then be compared to this random distribution. In other words, a PLV was significant if larger than 0.05 (noise floor) in our study. For all these analyses, a subset of 30 electrodes was chosen for analysis. These 30 electrodes were chosen as the electrodes with the largest amplitude of the onset response to the target and masker complex. These can be seen as the electrodes with maximum negativity as seen in the topographic maps (see Supplementary Figure 5). The PLVs were averaged across the subset of electrodes for each condition of each participant.

Target enhancement was calculated as the difference in PLV between the ENH and MSK conditions at the target AM frequency; masker enhancement was calculated as the difference in PLV between the ENH and MSK conditions at the masker AM frequency; and overall enhancement was calculated as the difference between the target and masker enhancement:

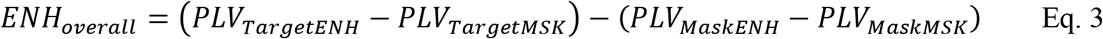

## ACKNOWLEDGEMENTS

This study was supported by NIH grant R01 DC012262 (AJO). We thank Patricia Leach and PuiYii Goh for assistance with data collection and Hao Lu for advice regarding EEG data analysis. Stephen Engel provided helpful comments on an earlier draft.

## Supplementary Figures

**Supplementary Figure S1:**
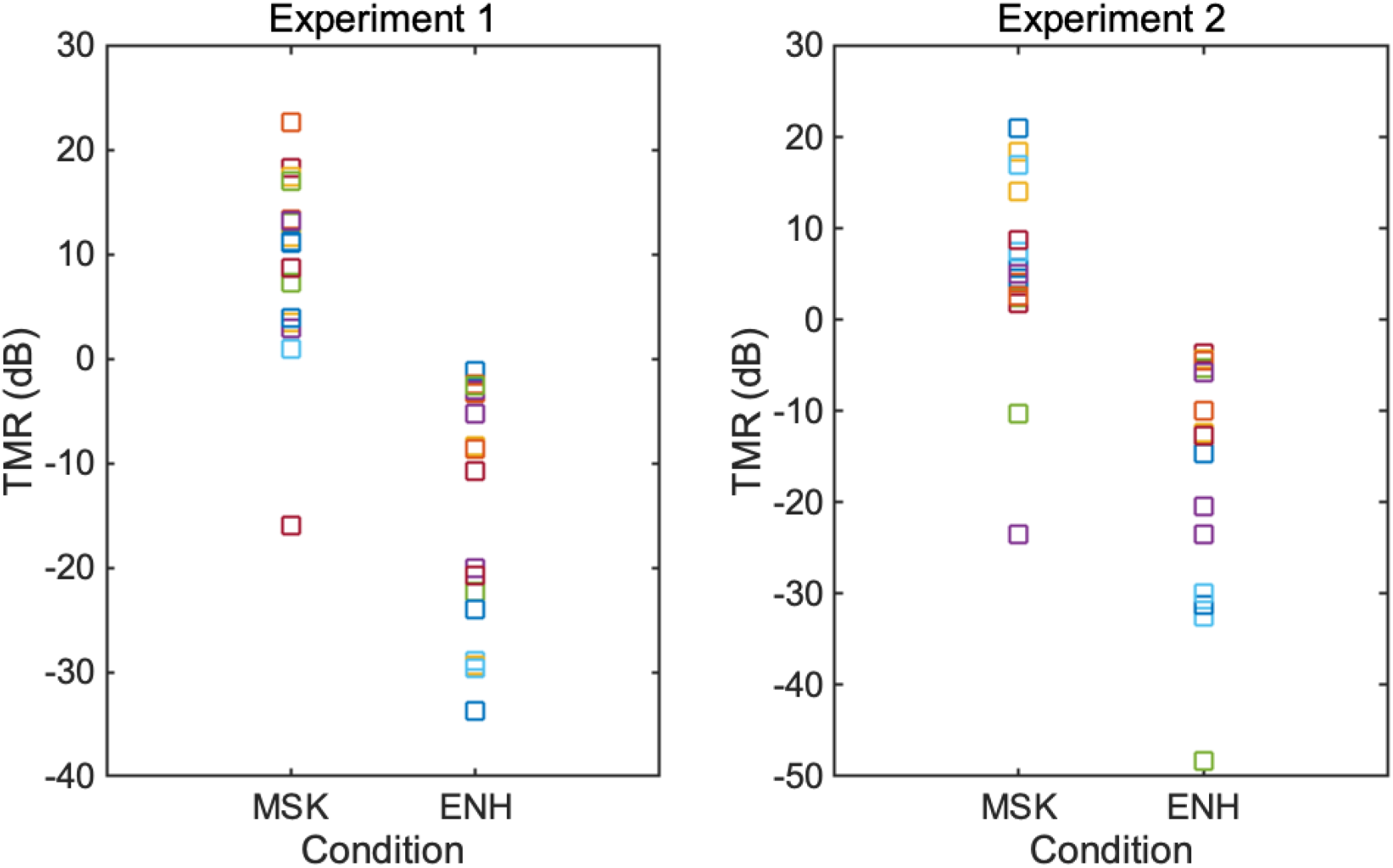
Individual behavioral thresholds from Experiments 1 and 2 of participants who passed the behavioral screening and were recruited for the EEG experiment.

**Supplementary Figure S2:**
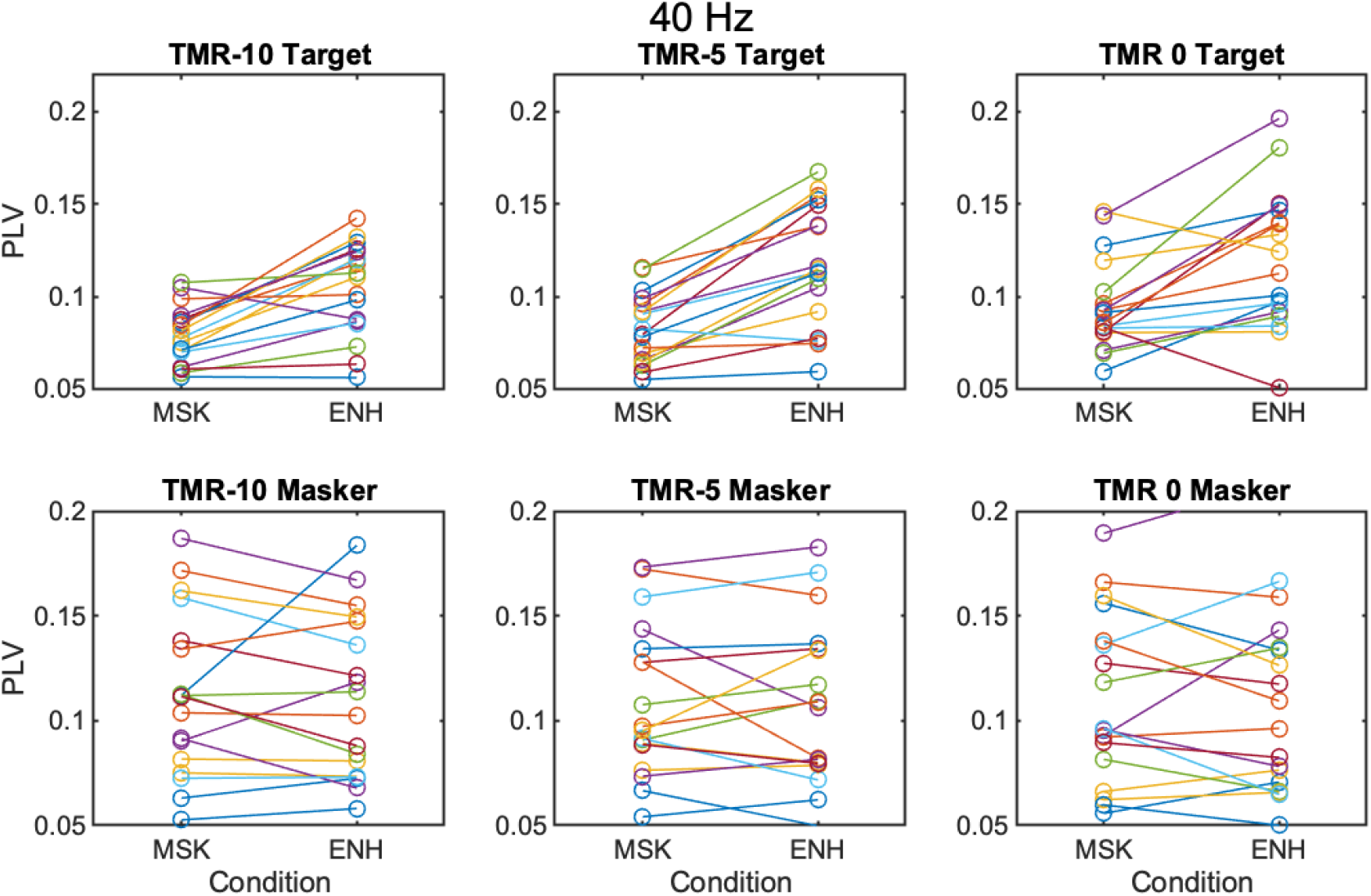
Individual PLV for each subject for Experiment 1 at the low modulation frequencies (~40 Hz). The three columns indicate the three TMR conditions. The top row indicates thresholds for the target modulation frequency and the bottom row indicates PLV for the masker modulation frequencies.

**Supplementary Figure S3:**
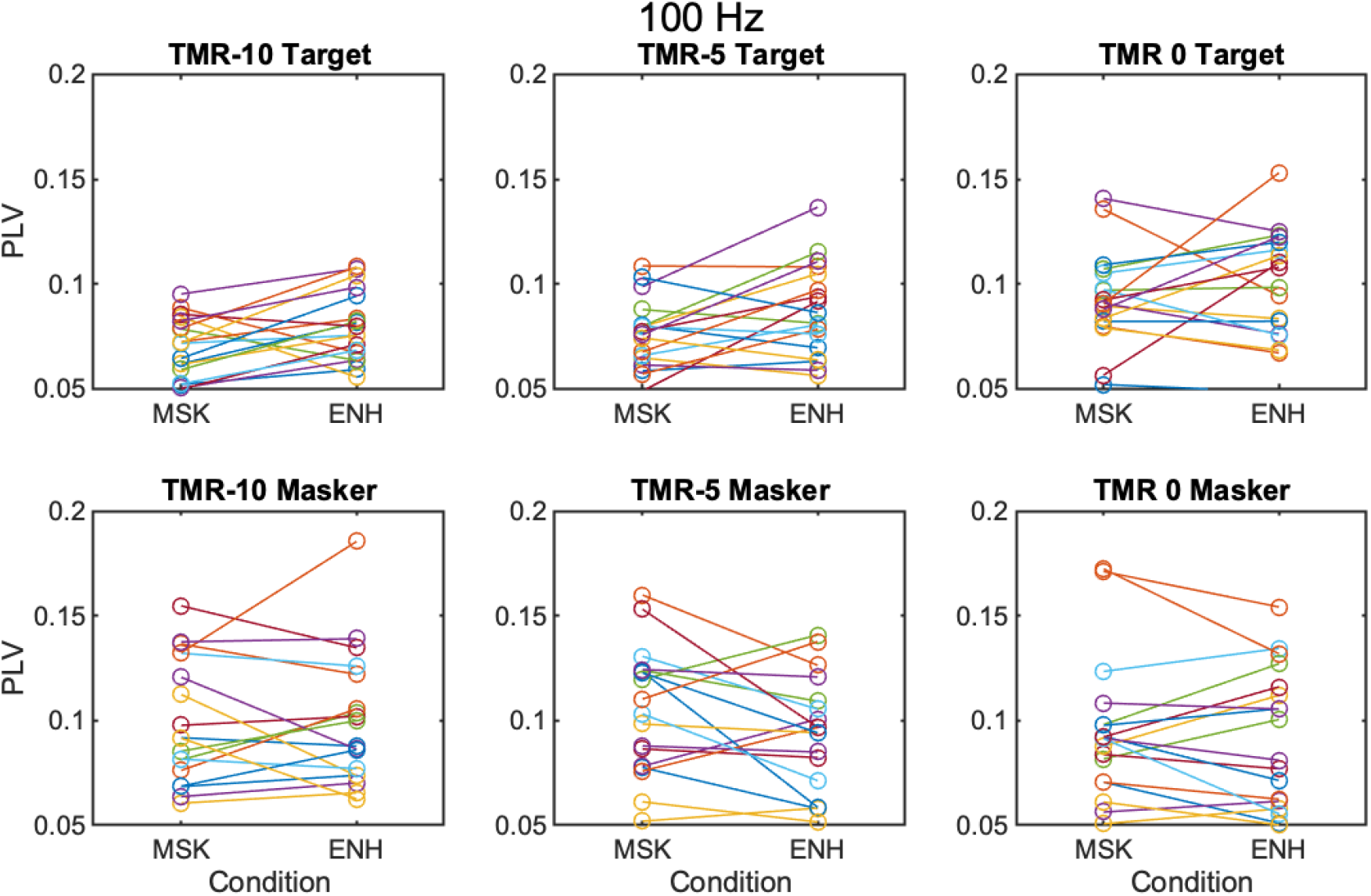
Individual PLV for each subject for Experiment 1 at the high modulation frequencies (~100 Hz). The three columns indicate the three TMR conditions. The top row indicates thresholds for the target modulation frequency and the bottom row indicates PLV for the masker modulation frequencies.

**Supplementary Figure S4:**
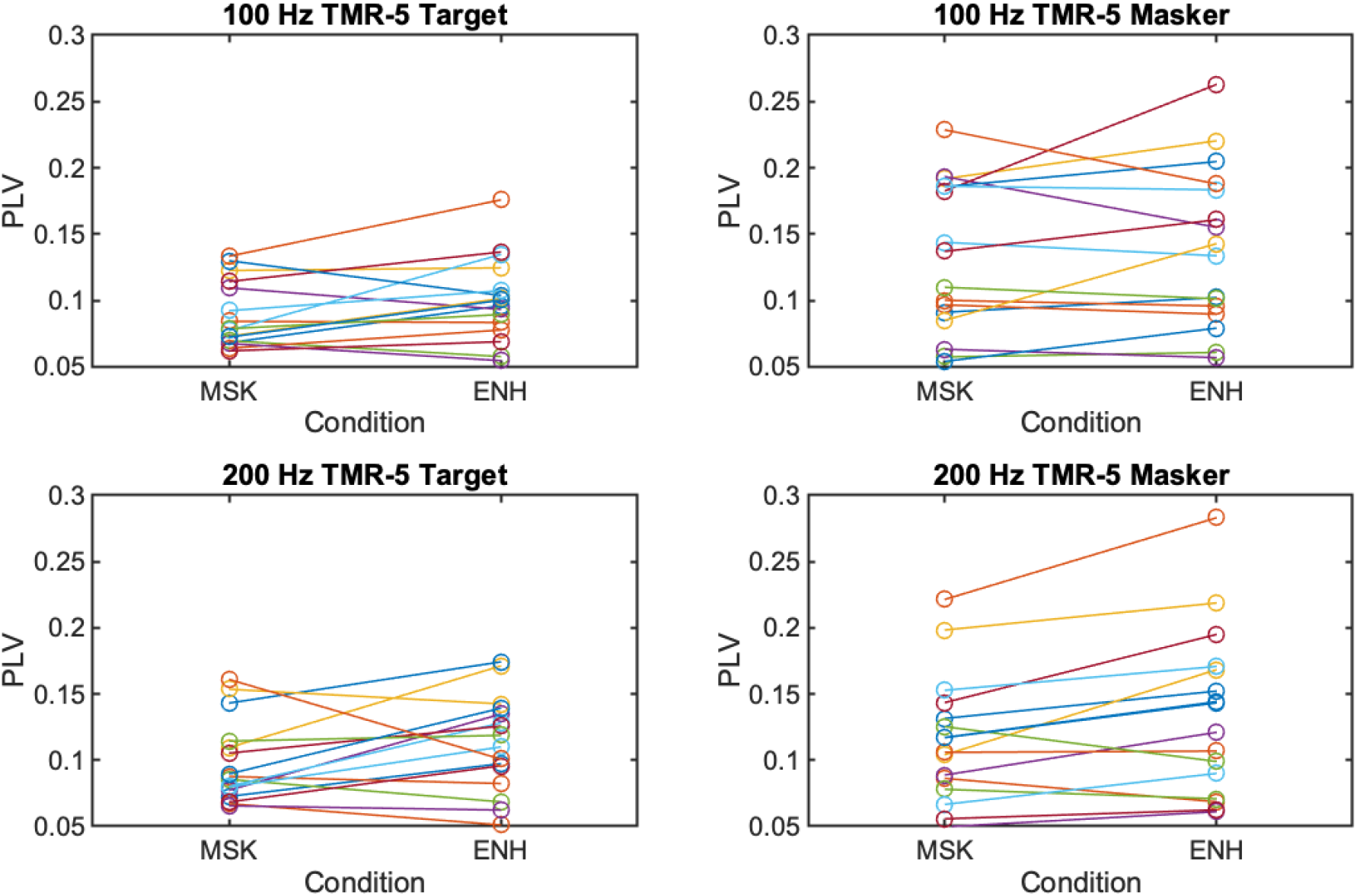
Individual PLV for each subject for Experiment 2. The top row indicates PLV for the ~100 Hz modulation frequencies and the bottom row indicates PLV for the ~200 Hz modulation frequencies. The first row is for the target modulation frequencies and the second row is for the masker modulation frequencies.

**Supplementary Figure S5:**
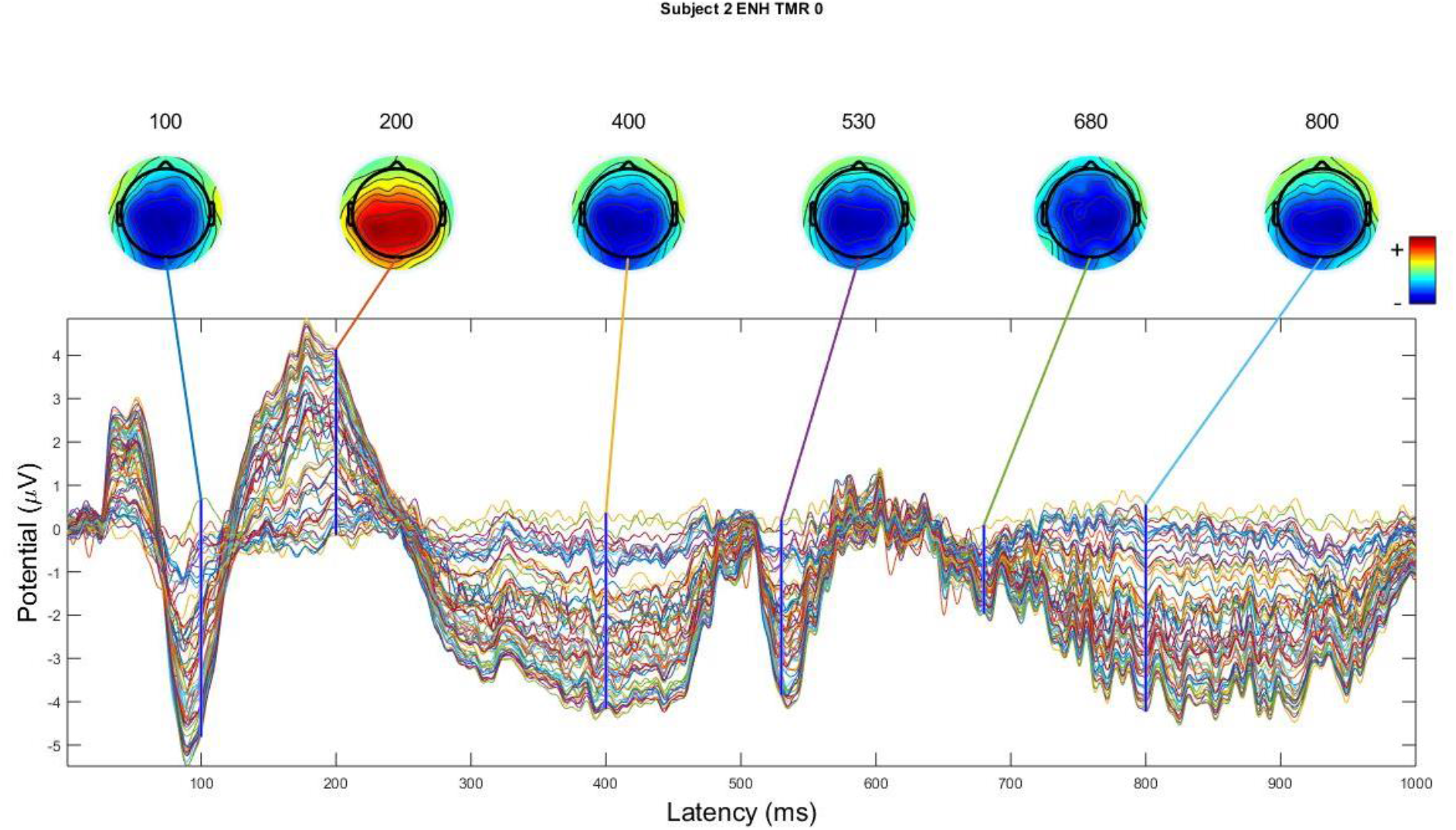
Example topographical plots (topoplots) for a sample subject at various time points in an averaged epoch for the ENH condition at TMR = 0 dB. Each waveform corresponds to a different electrode. The topoplots indicate the scalp distribution of the EEG activity.

**Supplementary Table 1.**
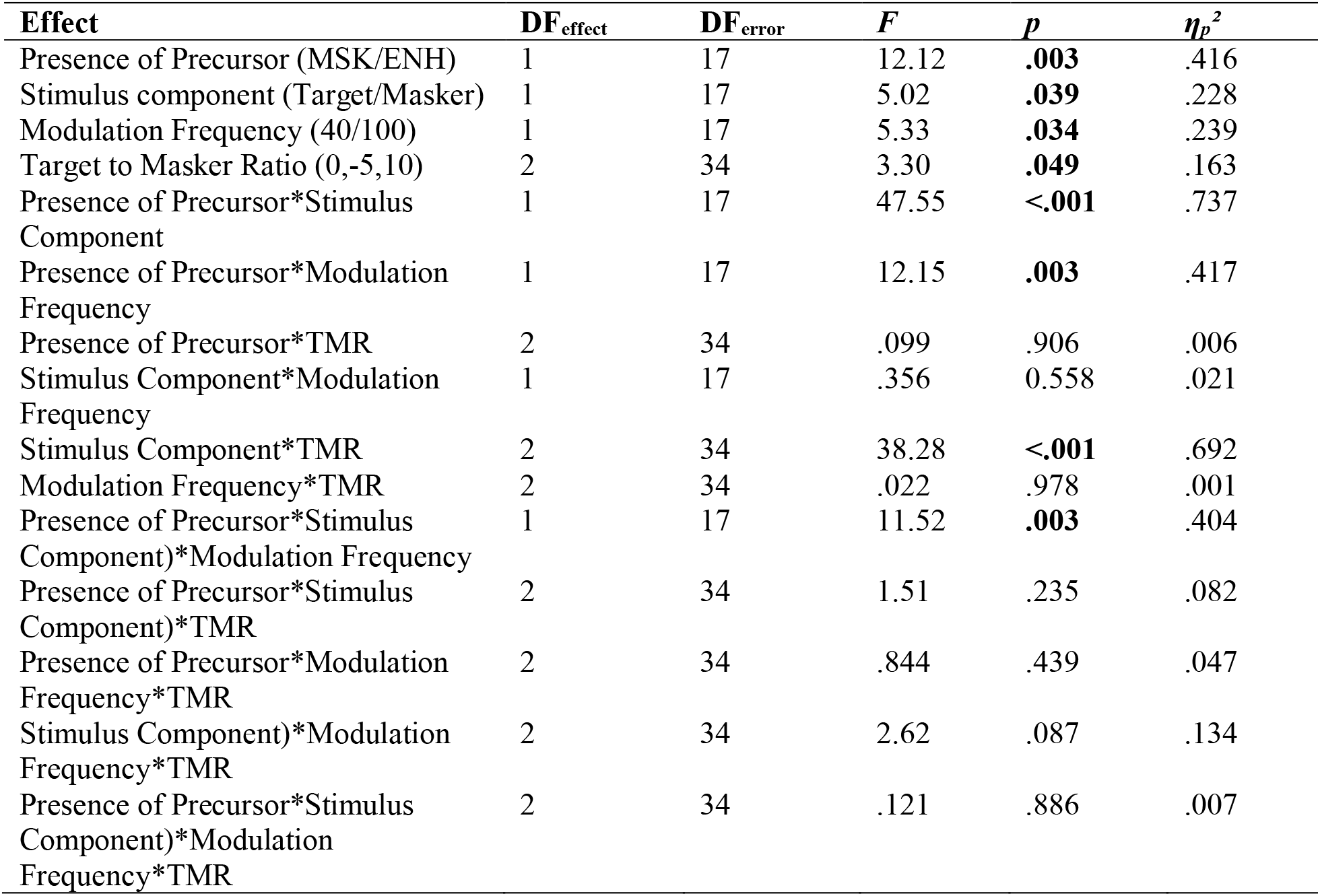
Experiment 1 results. Post hoc pairwise comparisons (with Bonferroni correction for multiple comparisons) showed that PLVs were significantly different between the MSK and ENH conditions for only the target components (for the low rate, p<0.001; for the high rate, p=0.034). PLVs were not significantly different for the maskers (for the low rate, p=0.469; for the high rate, p=0.223).

**Supplementary Table 2.** Experiment 2 results. No post-hoc tests were carried out as no interactions were significant.

| **Effect** | **DF<sub>effect</sub>** | **DF<sub>error</sub>** | **F** | **p** | **$\eta_p^2$** |
| --- | --- | --- | --- | --- | --- |
| Presence of Precursor (MSK/ENH) | 1 | 15 | 10.43 | **.006** | .410 |
| Stimulus component (Target/Masker) | 1 | 15 | 17.05 | **.001** | .532 |
| Modulation Frequency (100/200) | 1 | 15 | .000 | .987 | .000 |
| Presence of Precursor*Stimulus Component | 1 | 15 | .008 | .932 | .001 |
| Presence of Precursor*Modulation Frequency | 1 | 15 | 1.257 | .280 | .077 |
| Stimulus Component*Modulation Frequency | 1 | 15 | 2.473 | 0.137 | .142 |
| Presence of Precursor*Stimulus Component)*Modulation Frequency | 1 | 15 | .299 | .593 | .020 |

